# Stay tuned: active amplification tunes tree-cricket ears to track temperature-dependent song frequency

**DOI:** 10.1101/050583

**Authors:** Natasha Mhatre, Gerald Pollack, Andrew Mason

## Abstract

Tree cricket males produce tonal songs, used for mate-attraction and male-male interactions. Active mechanics tunes hearing to conspecific song frequency. However, tree cricket song frequency increases with temperature, presenting a problem for tuned listeners. We show that the actively amplified frequency increases with temperature, thus shifting mechanical and neuronal auditory tuning to maintain a match with conspecific song frequency. Active auditory processes are known from several taxa, but their adaptive function has rarely been demonstrated. We show that tree crickets harness active processes to ensure that auditory tuning remains matched to conspecific song frequency, despite changing environmental conditions and signal characteristics. Adaptive tuning allows tree crickets to selectively detect potential mates or rivals over large distances and is likely to bestow a strong selective advantage by reducing mate-finding effort and facilitating intermale interactions.

## Main text

Animals using acoustic signals encode considerable information about species identity and individual quality in song spectral and temporal features [reviewed in 1]. For receivers, however, separating conspecific signals from other sounds can be difficult in real-world conditions. Tree crickets signal in a crowded acoustic space with potential for masking[2]. Many insects deal with this problem through tuned auditory systems that respond selectively to frequencies representing conspecific signals[1]. However, in tree crickets this strategy is complicated by temperature-dependent, hence variable, song frequency[3].

Tree cricket auditory tuning is determined by frequency-specific, active amplification of tympanal vibration at low sound amplitudes[4,5]. Active amplification is a mechanical process that is achieved physiologically[6]; auditory neurons use axonemal dyneins[7,8] to generate forces[9] that preferentially amplify frequencies representing conspecific songs[4,5,10]. Since a physiological rather than mechanical process determines tuning, temperature is expected to affect this process.

Here we test, in the North American tree cricket *Oecanthus nigricornis*, whether the amplified frequency increases with temperature thereby allowing the auditory system to remain tuned by tracking changes in conspecific song frequency.

## Temperature affects self-sustained oscillations and mechanical tuning

Self-sustained oscillations (SOs) are characteristic of active auditory mechanics, and occur in tree cricket tympana[4,5]. SOs are generated by the same processes underlying active amplification, and reveal the actively amplified frequency[4,9,11]. Fig. 1A shows frequency spectra of SOs measured from the anterior tympanal membrane (ATM) over temperatures ranging from 21 to 30°C, over which SO frequency increased linearly by 112 Hz/°C (Fig. 1A inset). Change in SO frequency corresponded well with the previously reported range of *O. nigricornis* song frequencies (3.6 to 4.9 kHz from 19.3 to 29.6°C ambient temperature), and with the slope of the frequency-temperature relationship (144 Hz/°C, R^2^ = 0.74; data from [12]). The shallower slope that we measured may be due to a temperature mismatch between the surface of the Peltier element that we used to control temperature and the tympanum (see supplemental methods).

**Figure 1.**
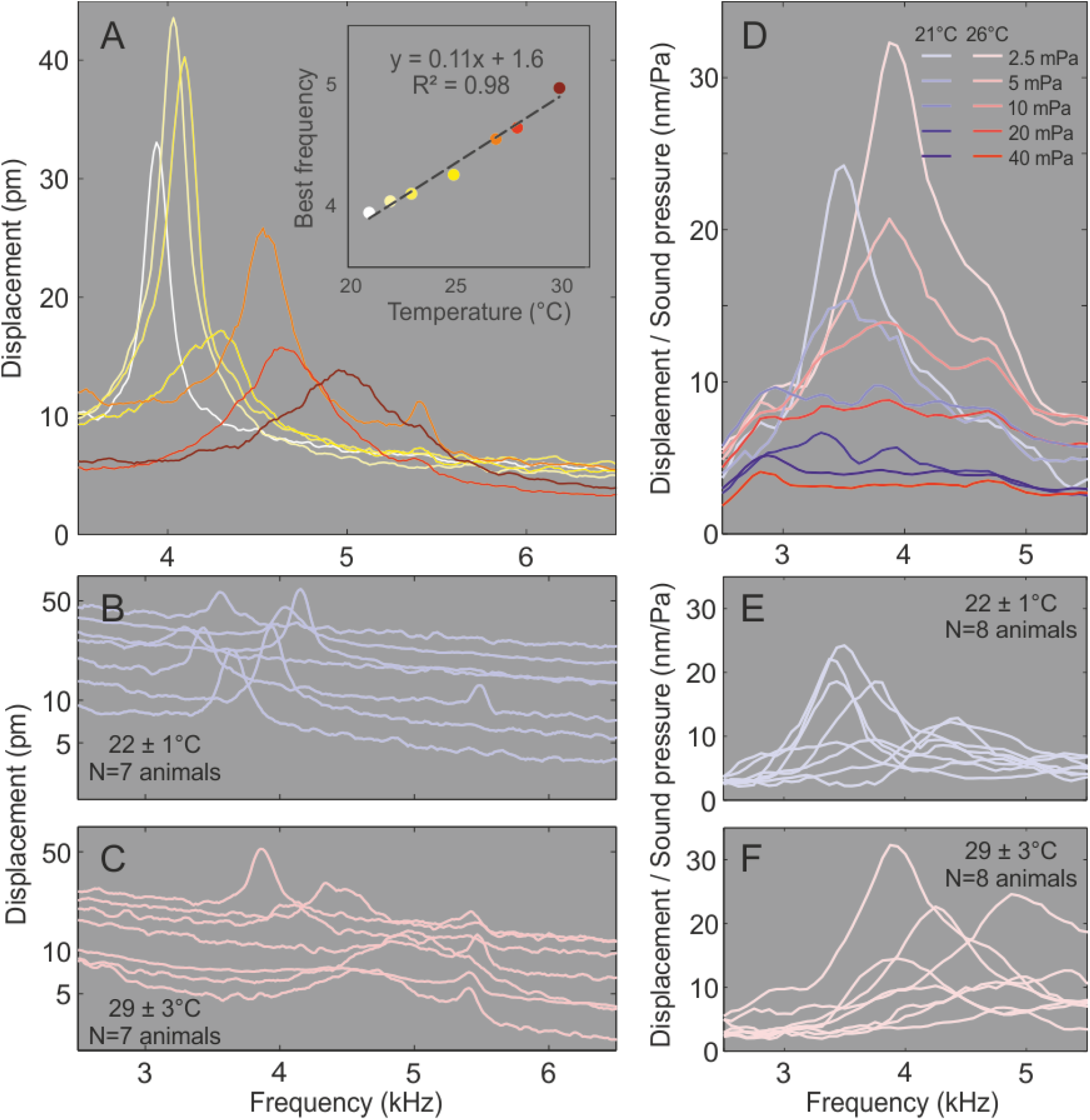
Spontaneous oscillations (SOs) and mechanical tuning of an example *Oecanthus nigricornis* ATM. (A) SO best frequency increases linearly with temperature. Colours in the inset correspond to the SO frequency spectra in the main plot. (B) SO frequencies at low temperature are lower than those at (C) high temperature. (D) Mechanical responses of ATM change with stimulus level and temperature. Mechanical tuning is sharpest at low stimulus amplitude and shifts to higher frequencies with increasing temperature. This shift in best frequency is consistent across individuals (E, F; stimulation level: 2.5 mPa).

Fig. 1B & C illustrate SO spectra for seven individuals (3 females, 4 males) at low (22±1°C) and high (29±3°C) temperatures. SO frequencies increased with temperature despite considerable variation in both temperature ranges (Table 1). In all seven individuals best frequency was higher at higher temperatures, increasing on average by 720 ± 470 Hz.

**Table 1.** Mechanical and neuronal tuning in different temperature regimes. Data are given as mean ± SD (N).

| | Low temperature<br>(22±1°C) | High temperature<br>(29±3°C) | Paired t-test |
| --- | --- | --- | --- |
| SO frequency (kHz) | 3.7 ± 0.3 (7) | 4.4 ± 0.4 (7) | * 1-tailed, P = 0.003 |
| Mechanical best frequency (kHz) | 3.8 ± 0.4 (8) | 4.4 ± 0.4 (8) | * 1-tailed, P = 2e-4 |
| Neuronal best frequency (kHz) | 3.3 ± 0.2 (8) | 4.3 ± 0.21 (8) | * 1-tailed, P=3.67e-5 |
| Q factor of mechanical response | 7.7 ± 3.6 (8) | 5.6 ± 1.8 (8) | 2-tailed, P=0.08 |

ATM motion showed the stimulus amplitude-dependent tuning well known from actively amplified auditory systems (Fig. 1D) [4,5]. At low stimulus amplitudes, a peak emerged in the ATM frequency response (Fig. 1D). Fig. 1 E & F show frequency response functions at low and high temperatures for 2.5 mPa (42 dB SPL) stimuli. As with the SOs, peak frequency was higher at higher temperatures in each individual. On average, best frequencies of sound-driven responses increased by 632 ± 270 Hz. The sharpness of the peak, (Q factor) was not significantly different between the two temperature ranges (Table 1).

## Temperature affects neural tuning

We recorded extracellularly from the prothoracic ganglion, the first site of central neuronal processing of auditory information in crickets (Fig. 2A, B). Recodings were made in the hemiganglion contralateral to the side of stimulation in eight *O. nigricornis* (4 of each sex). Four animals had only one ear, because of injury either during development or dissection. In field crickets, the only prothoracic neuron readily recorded contralateral to the site of afferent input is ON1 (see supplemental methods), whose frequency tuning is representative of the auditory system [13]. Although prothoracic auditory neurons of *Oecanthus* have not yet been characterized, given the recording site, and the occurrence of an ON1-like neuron in several ensiferans[13–15], it is likely that these recordings were from a neuron homologous to ON1.

**Figure 2.**
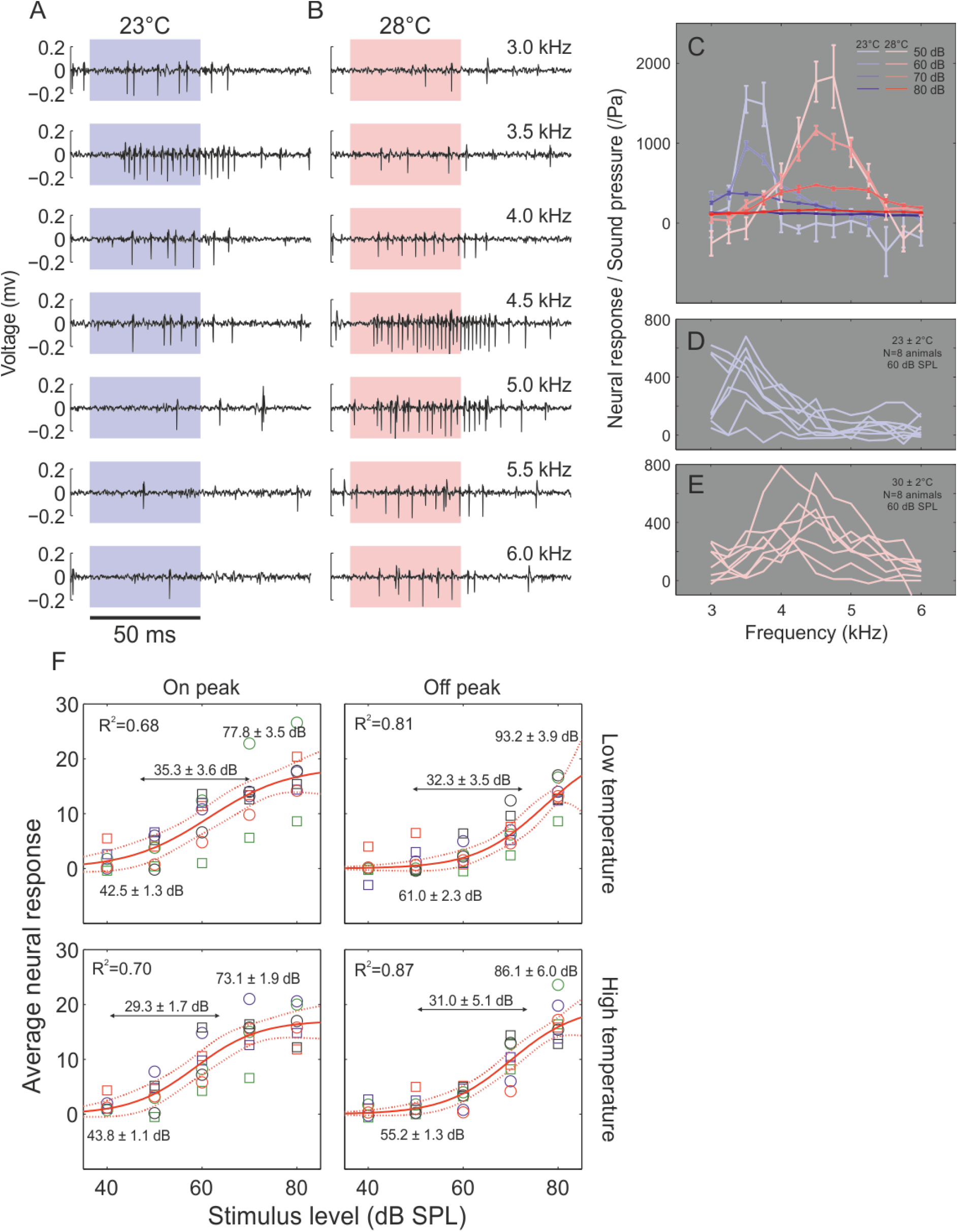
Sound-evoked responses of *O. nigricornis* putative ON1 interneuron at different temperatures. Example responses of one individual; 60 dB SPL stimuli at different frequencies for (A) low and (B) high temperatures. Coloured boxes indicate stimulus duration. (C) Tuning of an individual, expressed as sensitivity (neural response per unit sound pressure) varies with stimulus amplitude and temperature. Increases in best frequency with temperature are consistent across animals (D, E). (F) Thresholds are lower for on-peak frequencies than for off-peak frequencies (n = 8). Error lines indicate 95% prediction intervals for logistic fits. Threshold, saturation and dynamic range (mean ± SD) are indicated; variation is estimated by jack-knife resampling. Colours represent different individuals; males are indicated by circles and females by squares.

To allow comparison with mechanics, neural responses are represented per unit sound pressure (Fig. 2C–E). As observed in the mechanics, both neuronal tuning and sensitivity were amplitude-dependent. Tuning to conspecific frequency was sharpest near threshold and broadened as amplitude increased (Fig. 2C). Note, however, that amplitude-dependent tuning at the neural level does not require active auditory mechanics. Variation among receptors in sensitivity and tuning, and response saturation at high stimulus level, can account for responses that are more frequency-selective at low than at high stimulus levels. Indeed, we observed this in ON1 responses in the field cricket *Gryllus assimilis*, a species in which there is no sign of active tympanal mechanics (Fig. S1).

Crucially, however, neural tuning in *O. nigricornis* shifted up with temperature, mirroring the mechanics (Fig. 2D, E, Table 1). The average per-individual shift in best frequency was 938 ± 320 Hz. No such tuning shift occurred in *G. assimilis* (Fig. S1), reinforcing the conclusion that the shift in frequency tuning is implemented by active mechanisms.

## The benefit of temperature dependent tuning

Without active tuning, how much would a temperature mismatch reduce male active acoustic space? To estimate this, we compared the neuronal response at the best frequency at the temperature at which the recording was made (on-peak) and best frequency in the other temperature range (off-peak). Thresholds were determined from logistic fits to neuronal responses (Fig. 2F).

At both temperatures, thresholds were much lower for the amplified frequency (Fig. 2F, low temperature: P = 1.4e-4; high temperature: P = 4e-4; 1-tailed paired t-tests). Based on the shift in threshold, with fixed tuning to the low-temperature frequency, song produced at the best frequency for high temperatures would have to be 18.5 ± 1.8 dB SPL louder to be detected by a low-temperature ear. Considering attenuation due to spherical spreading alone, active amplification increases detection distance by a factor of 8.6 ± 1.7 and active area by a factor of 76.0 ± 30.9. Similarly song at best auditory frequency for low temperature would have to be 11.4 ± 1.8 dB SPL louder to be heard by a warm ear. This conservatively corresponds to an increase in detection distance by a factor of 3.8 ± 0.8 and acoustically active area by a factor of 14.8 ± 6.3.

The main effect of active amplification, thus, was to lower auditory thresholds at the temperature-appropriate frequency thereby increasing the distance over which a male signalling at that frequency could be heard.

It is important to note that the threshold achieved by the actively amplified *O. nigricornis* auditory system, at about 45 dB SPL, is similar to ON1 thresholds in field crickets ([13,16,17]). Indeed, even at very highest observed amplification, *O. nigricornis* tympana rarely achieve mechanical sensitivity higher than *G. assimilis* (Fig 1, S1, [5]). It has been argued that an actively amplified auditory system produces a high level of spontaneous noise near criticality, and so cannot lower the minimal detectable sound level[18]. Thus, we find that the primary function of active amplification in tree-crickets is not to lower the absolute auditory threshold below that of other crickets but rather to modulate auditory tuning to match conspecific song frequency even as it changes with temperature.

Active amplification has been invoked to explain the remarkable sensitivity, dynamic range, and frequency resolution of many auditory systems[19]. However, few studies demonstrate the contribution of active amplification to these processes, especially in ecologically relevant settings. Given that some features, such as fine frequency tuning[20] and high dynamic range[21] are achievable in systems that are, to the best of our knowledge, purely passive, the advantage of active amplification remains to be demonstrated.

Perhaps the most compelling demonstrations of functional roles for active amplification come from fruit flies and mosquitoes. In fly courtship, active amplification contributes to species isolation by matching antennal tuning to conspecific song frequency [10]. However, passive resonance can also achieve matched-filtering (Fig. S1 A, B, [22]), and the specific advantage of active tuning is unclear. In mosquitoes, mate recognition is believed to be mediated by convergence of higher harmonics of male and female flight tones[23]. In the genus *Culex*, mismatched harmonics are thought to be detected using low-frequency difference tones generated by active mechanics[24]. However, in *Aedes aegypti* mosquitoes, the frequencies of the higher harmonics can be detected directly by the auditory organ[25], again questioning the specific advantage of the active process.

We show that an important selective advantage of active amplification is its flexibility and responsiveness to environmental conditions. While sender-receiver match can be achieved through resonance for a fixed frequency signal[22], resonant systems can only remain sensitive to a variable-frequency signal by being broadly tuned, thus compromising selectivity. Active systems can modify the amplified frequency and bypass this trade-off.

In insects, active amplification depends on the action of ATP dependent molecular motors[26]. Models suggest that the amplified frequency, and hence tuning may be determined by the ATPase rate of these motors[7,27]. Indeed, the temperature dependence of SO frequency has been observed in two poikilothermic organisms, frogs[28] and mosquitoes[7]. Here we show that tree crickets have both temperature-dependent SOs and mechanical and neural tuning, and that they exploit this property to maintain auditory sensitivity to song even as it changes with temperature, without loss of selectivity.

## Data accessibility

The dataset supporting this article has been uploaded to the Dryad repository: https://datadryad.org/resource/doi:10.5061/dryad.fv8jv

## Author contributions

NM made mechanical measurements and analysed data, GP made neuronal measurements and analysed data, NM, GP and AM designed the study and wrote the manuscript. All authors agree to be held accountable for the content therein and approve the final version of the manuscript.

## Acknowledgements

This research was funded by NSERC Discovery grants to Gerald Pollack and Andrew Mason.

## Supplementary materials and methods

### Animals

*Oecanthus nigricornis* were collected from fields in Scarborough, Ontario, Canada. Animals were maintained collectively in a large cage with 12h:12h day:night conditions with *ad libitum* access to food and water.

### Vibrometry

Animals were fixed in position on a steel rod with a rotatable platform and positioned in the path of the laser using a previously described technique[1]. Animals were alive throughout the duration of the experiment as verified by breathing movements. Vibrometric measurements were made from the anterior tympanal membrane (ATM) and the data reported are from the point of maximal displacement[2,1,3]. Acoustic stimuli were 16 ms periodic chirps from 2 to 20 kHz with a frequency resolution of 62.5 Hz produced and re-acquired at a sampling rate of 128 kHz. Each stimulus type was presented 100 times and the response of the ATM was averaged across all presentations using the complex averaging procedure, which accounts for both the magnitude and phase of the response. The stimuli were produced by the vibrometer software (Polytec Scanning vibrometer software version 9.1.2) through a National instruments DAQ board (PCI 6110), amplified and broadcast via a loudspeaker. All acoustic stimulation was ipsilateral and was continuously monitored during the vibrometry experiments. A Bruel and Kajaer 1/4 inch microphone (Type 4231) (frequency response ± 1 dB: 20 Hz to 80 kHz) was placed 1 cm above the animal’s ATM and connected to a Bruel and Kajaer SPL meter (Type 2231), the A/C output from which was continuously monitored as a reference by the vibrometer software. The delivered sound stimulus was flat (± 3 dB) across the frequency range of interest. The average FFT amplitudes of the delivered stimuli were 2.5, 5, 10, 20 and 40 mPa. These correspond to peak or instantaneous amplitudes of 50, 100, 200, 400 and 800 mPa respectively.

Vibration measurements were performed with a Polytec (PSV 400) scanning laser-vibrometer coupled with an OFV 505 sensor fitted with a close-up unit and acquired through the same NI DAQ PC 6110 board used for stimulus generation. The entire experimental apparatus was placed on a vibration isolation table (Newport VH3036 W-OPT, Irvine, CA, USA) in an acoustic isolation booth (Eckel XHD-BATTEN, ON, Canada). The vibrometer measured the velocity of the ATM as it vibrated in response to acoustic stimulation, using the same FFT parameters as used for stimulation. Given the low amplitude of the ATM vibrations, the lowest range velocity decoder available for vibration detection was used (VD 09 5 mm/s/V). A transfer function between the acoustic stimulation and vibrational response was then calculated[4] allowing us to relate the amplitude and phase of the displacement of the ATM to the amplitude and phase of the acoustic stimulus. The data are plotted after being smoothed over a 5 point span using a moving average. The resonant peak in the transfer function (at 2.5mPa stimulation level) was fitted to a simple harmonic oscillator model and the Q factor calculated from the damping ratio [1]. The ATM was also observed to oscillate spontaneously in the absence of sound. Oscillations were recorded using the same decoder at a sampling rate of 128 kHz and their frequency spectra were calculated from 2 to 20 kHz at a resolution of 15.6 Hz. These data are plotted after being smoothed over an 11 point span using a Savitzky-Golay smoothing algorithm.

### Temperature control

Vibrometric measurements were made at room temperature (22±1°C) and at high temperatures (29±3°C).The animal was glued to the surface of a 40 × 40 mm Peltier plate, the surface temperature of which was controlled by a variable voltage and monitored using a thermocouple coupled with a voltmeter. We report the surface temperature of the Peltier plate. Control measurements showed that with plate temperature at 30°C, temperature in the air 2 mm above the surface, the approximate position of the ear during measurements, was 28°C. During measurements at high temperature, ear temperature is likely to be somewhat warmer than surrounding air due to heat conduction within the cricket’s body; thus we estimate that plate temperature may overestimate ear temperature by less than 2°C.

### Electrophysiology

After removing the meso- and metathoracic legs and the wings, animals were mounted ventral side uppermost on a 15 mm × 15 mm Peltier plate such that the dorsal surfaces of the thorax and abdomen were in contact with the plate. The prothoracic coxae were retracted from the midline with small hooks, the legs were flexed at the femor-tibial joint in a nearly natural position, and the tarsi were waxed to vertical pins. Temperature of the Peltier plate was controlled and measured as described above.

The prothoracic ganglion was exposed by ventral dissection, supported on a small metal platform and bathed in physiological saline[5]. Tungsten microelectrodes with resistance of approximately 5 MOhm recorded from sound-activated units in the hemiganglion opposite to the side of stimulation. In most recordings, responses of single units could be identified (see Fig. 2). In two animals, the prothoracic leg, and thus the ear ipsilateral to the recording electrode was autotomized during preparation for recordings. In a third animal the electrode-ipsilateral leg was removed deliberately following the measurements of frequency sensitivity. In field crickets, the well characterized ON1 neuron receives afferent input on dendritic processes in one hemiganglion, and provides output via axonal processes in the other hemiganglion. The only known prothoracic neuron that can be recorded reliably (from its axonal processes) contralateral to the intact ear in the one-eared animals described above is ON1. Following removal of one ear, the deafferented ON1 can re-acquire afferent input from the remaining ear by producing dendritic sprouts that cross the midline[6]. The brief time between leg removal and recording, however, precludes this possibility for the three animals described above. A fourth animal lost most of its prothoracic leg (as an adult, as indicated by scarring) in the wild before collection; thus dendritic sprouting cannot be ruled out in this case.

Stimuli were 50 ms tones, including 1 ms onset and offset ramps, presented once per second with frequencies from 3 to 6 kHz in steps of 0.25 kHz, at sound levels from 40 or 50 to 80 dB SPL in steps of 10 dB. The loudspeaker was at an angle of 45° from the midline at a distance of 21 cm. The entire set of stimuli was presented five times, and responses of each cricket were averaged over the five repetitions. To account for spontaneous activity, the neural response to each stimulus was defined as the difference in number of spikes (as detected by thresholding) in the 100 ms preceding and following stimulus onset.

Neural responses were fitted with a logistic function (Fig. 2F), where, the response (y) was modelled as a function of the stimulus level (x) in dB SPL.

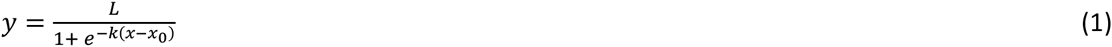

L is the response at the saturation asymptote and the x_0_ is the response in the middle of the dynamic range. Threshold level is defined as 10% of L and saturation as 90%. The dynamic range is defined as the difference in the stimulus level between threshold and saturation.

The logistic function was first fitted to the neural response pooled across all animals; this is the fit and the prediction intervals that are plotted in Fig. 3. To estimate the variance in the threshold and saturation, we jack-knife resampled the data. In a series of fitting exercises, each individual was dropped one by one and a new fit made to the truncated data set. Threshold and saturation were estimated for each fit and the means and standard deviations are reported.

**Figure S1.**
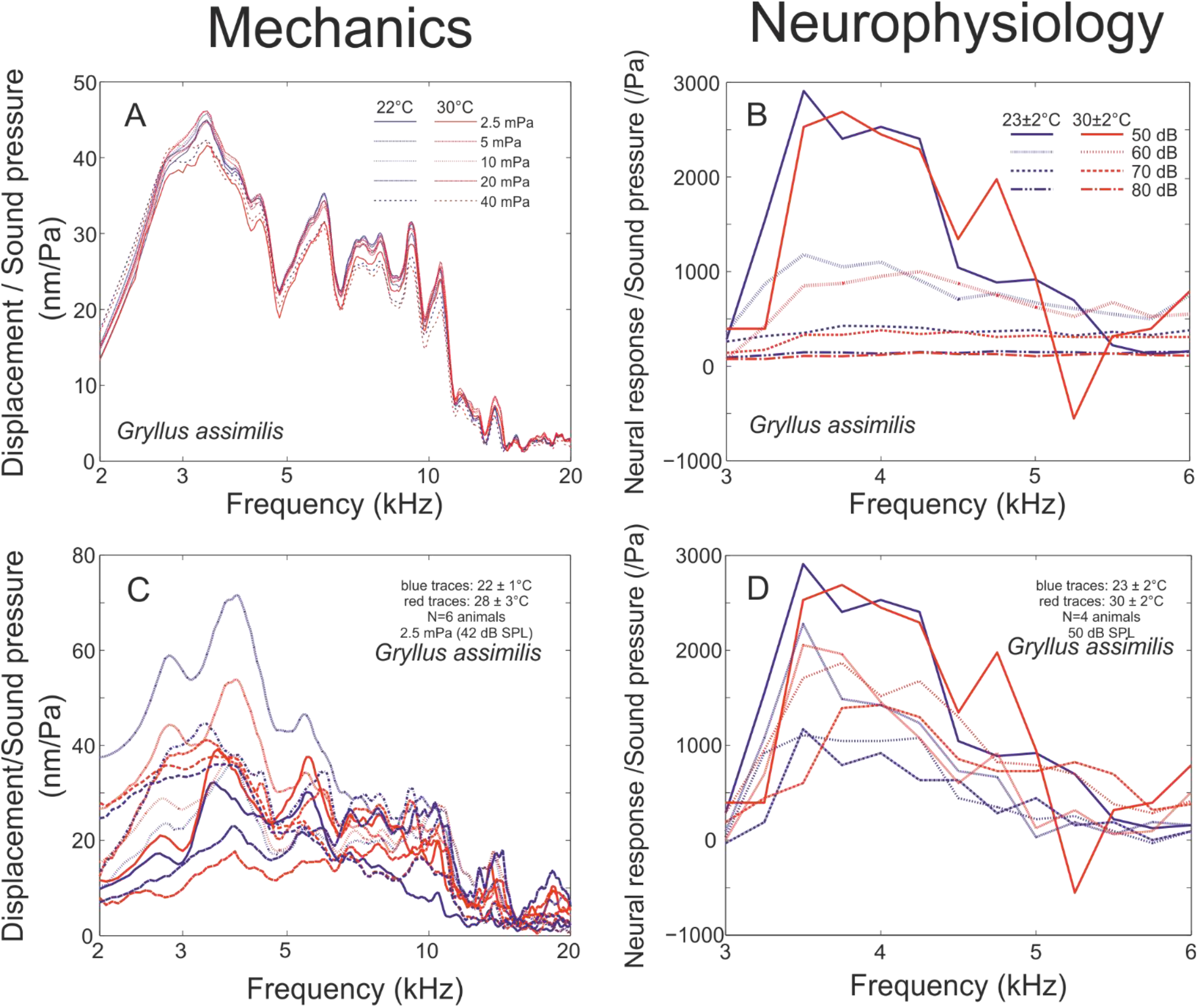
The mechanical and neurophysiological responses of *Gryllus assimilis*. (A) The mechanical tuning of the posterior tympanal membrane (PTM; the only functional tympanum of field-cricket ears) is independent of both stimulus amplitude and temperature. (B) The tuning of an example ON1 does not change with temperature, although sensitivity (neural response per unit sound pressure) decreases and becomes more broadly tuned as stimulus amplitude increases. At low stimulus amplitudes, it is clear that neither (C) the mechanical nor the (D) neuronal tuning of the auditory system changes with temperature.

